# Socio-sexual environment influences fecundity, but not response to bacterial infection, in *Drosophila melanogaster* females

**DOI:** 10.1101/2025.10.15.682585

**Authors:** Suhaas Sehgal, Aabeer Basu, Nagaraj Guru Prasad

**Affiliations:** Department of Biological Sciences, Indian Institute of Science Education and Research Mohali, India

**Keywords:** cost of immunity, female-female competition, mate harm, nonreciprocal trade-off, post-infection survival, reproduction-immunity trade-off

## Abstract

It is common for a host exposed to a pathogenic infection challenge to exhibit suppression of fecundity. This suppression can be driven by either a resource allocation-based trade-off between fecundity and immune function, or by infection-induced damage to host soma (particularly the reproductive organs). Alternatively, hosts can increase their fecundity to compensate for the curtailed lifespan resulting from lethal infection. Various factors, both extrinsic and intrinsic to the host, determine the actual effect of any infection challenge on host fecundity. While the host’s physiological and abiotic environmental factors have been thoroughly explored in this context, the role of the host’s biotic environment – especially its social and sexual environment – is rarely addressed. We investigated whether the socio-sexual environment (SSE) of *D. melanogaster* females influenced the effect of bacterial infection on their fecundity. We altered the socio-sexual environment of the females by housing them with different numbers of same-sex and opposite-sex companions. Our results indicate that while such alterations of the SSE changed the baseline reproductive effort of the females, they had no role in determining female susceptibility to infections or how infection affected female fecundity.

## 1. Introduction

Hosts challenged with a pathogenic infection commonly exhibit a reduction in their reproductive output (Nystrand and Dowling 2020). This reduction in fecundity is a manifestation of the virulence of the infection (Abbate et al., 2015). This reduction may be driven by a resource (re)allocation-based trade-off between reproduction and immune function (Sheldon and Verhulst 1996, Lochmiller and Deerenberg 2000, Sadd and Schmid-Hempel 2009): an infected host is expected to prioritise investment towards somatic maintenance and survival (Minchella 1985). The reduction in fecundity may also be driven by infection-induced damage to the host soma, particularly the reproductive organs, resulting from either the actions of the pathogen or those of the host immune response (Brandt and Schneider 2007, Sadd and Siva-Jothy 2006). An alternative possibility is that an infected host increases its immediate reproductive effort to compensate for the potential loss of life (and therefore future opportunities of reproduction) due to the pathogenic infection (Minchella and Loverde 1981, Parker et al., 2011).

The exact effect that a pathogenic infection challenge has on host fecundity is determined by various factors, both intrinsic and extrinsic to the host (Duffield et al., 2017). These factors either directly modulate the effect of the infection challenge on host reproduction, or influence it indirectly by affecting the risk of mortality from, and the degree of pathogenicity of, the infection (Sandland and Minchella 2003, Lazzaro and Little 2009). The roles of host physiological parameters (e.g., host age) and the abiotic components of the host environment (e.g., availability of resources and nutrition) have been explored thoroughly in this context (reviewed in Duffield et al., 2017). What is commonly overlooked is that the biotic components of the host environment, such as interactions with conspecific individuals, can also play a role in shaping how an infection challenge affects host fecundity. In this study, therefore, we explored the role of the host’s social and sexual environment in determining how infection affects host fecundity.

We used *Drosophila melanogaster* females infected with a bacterial pathogen as the model system for our investigations. Previous studies have demonstrated that the effect of bacterial infection on female fecundity in *D. melanogaster* can range from significant fecundity suppression to increased reproductive output, depending on various factors including (but not limited to) the identity of the infecting pathogen, host genotype and diet, and the phase of infection (Brandt and Schneider 2007, McKean et al., 2008, Linder and Promislow 2009, Howick and Lazzaro 2014, Kutzer and Armitage 2016, Kutzer et al., 2018, Kutzer et al., 2019, Hudson et al., 2020, Basu et al., 2024a).

*D. melanogaster* is neither a social insect nor a species that lives together in large numbers (Markow 2015). However, during certain life stages (e.g., larvae developing in a single fruit) and under specific ecological conditions (e.g., near oviposition sites), members of this species aggregate in substantial numbers (Reaume and Sokolowski 2006). These aggregations provide ample opportunities for inter-individual interactions of various natures, and such interactions can shape the behaviour and life history of these flies (Reaume and Sokolowski 2006, Billeter et al., 2025). There is sufficient evidence that social isolation and housing in different densities (ranging from small and benign changes in density to severely adverse and stressful conditions) can influence the physiology of *D. melanogaster* flies (Chen and Sokolowski 2022, Vora et al., 2022, Billeter et al., 2025). Importantly, changes in the socio-sexual environment have been previously shown to influence female reproductive behaviour and fecundity in *D. melanogaster* females (Fowler et al., 2022). Therefore, *D. melanogaster* flies are a suitable system for testing the effects of transient changes in social and sexual environments on responses to infections, both in terms of survival and fecundity.

We infected female flies with the bacterial pathogen *Enterococcus faecalis*, a known natural pathogen of this species (Lazzaro et al., 2006), and exposed these flies (along with sham-infected controls) to different socio-sexual environments. We manipulated the social and sexual environment independently by varying the number of female and male companions with which the focal females were housed, respectively. We aimed to test if altering the socio-sexual environment of the focal females determined their (a) fecundity, (b) post-infection survival, and (c) the effect of infection on their fecundity. Our results indicate that while the socio-sexual environment of the females influenced their fecundity, it did not affect their susceptibility to infection or the infection-induced suppression of fecundity.

## 2. Materials and methods

### 2.1. Pathogen handling and infection protocol

The Gram-positive bacterium *Enterococcus faecalis* (Lazzaro et al., 2006) was used in the experiments reported here. The bacteria are preserved as glycerol stock at −80 ^O^C. To obtain live bacterial cells for infections, 10 mL of lysogeny broth (Luria-Bertani Broth, Miler, HiMedia) is inoculated with the bacterial stock and incubated overnight with aeration (150 rpm shaker incubator) at 37 °C. 100 microliters of this primary culture are inoculated into 10 mL of fresh lysogeny broth and incubated for the necessary amount of time to obtain confluent cultures (OD600 = 1.0-1.2). The bacterial cells are then pelleted down using centrifugation and resuspended in sterile MgSO_4_ (10 mM) buffer at an optical density (OD_600_) of 1.0; OD_600_ = 1.0 for *E. faecalis* corresponds to 10^7^ cells/mL. Flies are infected, under light CO_2_ anaesthesia, by pricking the dorsolateral side of the thorax with a 0.1 mm Minutien pin (Fine Scientific Tools, CA, USA) dipped in the bacterial suspension. Sham-infections (controls for injury) are carried out in the same fashion, except by dipping the pins in sterile MgSO_4_ (10 mM) buffer.

### 2.2. Host population and generation of experimental flies

Flies from the <u>B</u>lue <u>R</u>idge <u>B</u>aseline <u>2</u> (BRB2) population – a large, lab-adapted, outbred *Drosophila melanogaster* population – were used as the focal flies in the experiments reported here. This population was established by hybridising 19 iso-female lines, which were founded from wild-caught females (Singh et al., 2015). The BRB2 population is maintained as a large, outbred population with a generation time of 14 days, comprising approximately 2,800 adults in each generation. Every generation, eggs are collected from the population cage (plexiglass cage: 25 cm length × 20 cm width × 15 cm height) and dispensed into vials (25 mm diameter × 90 mm height) with 8 mL banana-jaggery-yeast food medium at a density of 70 eggs per vial. 40 such vials are set up; the day of egg collection is demarcated as day 1. These vials are incubated at 25 °C, 50–60% RH, 12:12 hour LD cycle; under these conditions, the egg-to-adult development time for these flies is about 9–10 days. On day 12 post egg collection, all adults are transferred to a population cage and provided with fresh food plates (banana-jaggery-yeast food medium in a 90 mm Petri plate) supplemented with *ad libitum* live yeast paste. On day 14, a fresh food plate is provided in the cage, and 18 hours later, eggs are collected from this plate to initiate the next generation. All flies of the BRB2 population have the wild-type red eye colour.

Flies from the BL_st_ population (Dasgupta et al., 2019) – another large, lab-adapted, outbred *D. melanogaster* population – were used as the companion flies in experiment 1 reported here. The maintenance regime of the BL_st_ population is identical to that of the BRB2 population, except that the BL_st_ population is maintained under constant light. All flies of the BL_st_ population are homozygous for the recessive scarlet eye colour (Dasgupta et al., 2019).

For each replicate of the experiments, eggs were collected from the BRB2 population cage and distributed into food vials containing 8 mL of standard food medium at a density of 70 eggs per vial. These vials were incubated as per the general maintenance regime. On day 10 post egg-collection, freshly eclosing adults were collected as virgins within 6 hours of eclosion and housed in same-sex vials (10 females/vial or 12 males/vial). Virgin BL_st_ females were also obtained similarly for experiment 1 and housed in the same-sex vials (10 females/vial). To obtain inseminated females for experiments 1 and 2, on day 12 post egg-collection, virgin BRB females and males were combined in fresh food vials (in a 10:12 female-to-male ratio) and allowed to mate. These flies were housed in these vials till the time of infection (on day 14). Virgin BL_st_ females (experiment 1) and excess virgin BRB2 males (experiment 2) were transferred to fresh food vials on day 12 and housed therein till day 14. On day 14 post-egg collection, inseminated BRB2 females (along with the males) were transferred into fresh food vials 6 hours before infection.

### 2.3. Experiment 1: Effect of varying numbers of female companions on post-infection survival and fecundity of focal females

Inseminated BRB2 females were anaesthetised and subjected to infection (n = 60 per housing treatment) or sham infection (n = 30 per housing treatment); these females are hereafter referred to as the focal females. Post infection, focal females were housed in fresh food vials (with 8 mL of banana-jaggery-yeast food medium), either (a) individually, (b) with 1 companion BL_st_ female, or (c) with 3 companion BL_st_ females. All the companion BL_st_ females were virgins. The focal females were allowed to lay eggs in these vials for 48 hours, during which their survival was monitored every hour. For vials in which the focal female perished, the companion females (if present) were discarded, and the carcass of the focal female was removed from the vial. For vials in which the focal female did not perish by the end of 48 hours, all surviving females were discarded at the end of this period. All vials were then incubated under standard maintenance conditions, and 12 days later, the number of adult progeny in each vial was counted. The count of adult progeny in each vial was considered as the measure of fecundity of the focal female in that vial.

### 2.4. Experiment 2: Effect of varying numbers of male companions on post-infection survival and fecundity of focal females

Inseminated BRB2 females were anaesthetised and subjected to infection (n = 60 per housing treatment) or sham infection (n = 30 per housing treatment); these females are hereafter referred to as the focal females. Post infection, focal females were housed in fresh food vials (with 8 mL of banana-jaggery-yeast food medium), either (a) individually, (b) with 1 companion BRB2 male, or (c) with 2 companion BRB2 males. All the companion BRB2 males were virgins at the beginning of the experiment. The focal females were allowed to lay eggs in these vials for 48 hours, during which their survival was monitored every hour. For vials in which the focal female perished, the companion males (if present) were discarded, and the carcass of the focal female was removed from the vial. For vials in which the focal female did not perish by the end of 48 hours, all surviving flies were discarded at the end of this period. All vials were then incubated under standard maintenance conditions, and 12 days later, the number of adult progeny in each vial was counted. The count of adult progeny in each vial was considered as the measure of fecundity of the focal female in that vial.

### 2.5. Statistical analyses

All analyses were carried out using R statistical software, version 4.1.0 (R Core Team, 2021). Survival of infected females from both experiments was analysed using mixed-effects Cox proportional hazards models with ‘companion count’ as a fixed factor and ‘replicate’ as a random factor. Only data from infected females were included in the analyses, as sham-infected females did not exhibit any mortality within the observation window in either experiment (**figure 1**). Female fecundity (progeny produced per hour) in each vial was normalised before analysis as follows:

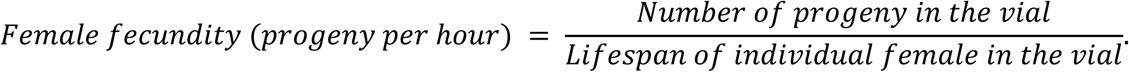

**Figure 1.**
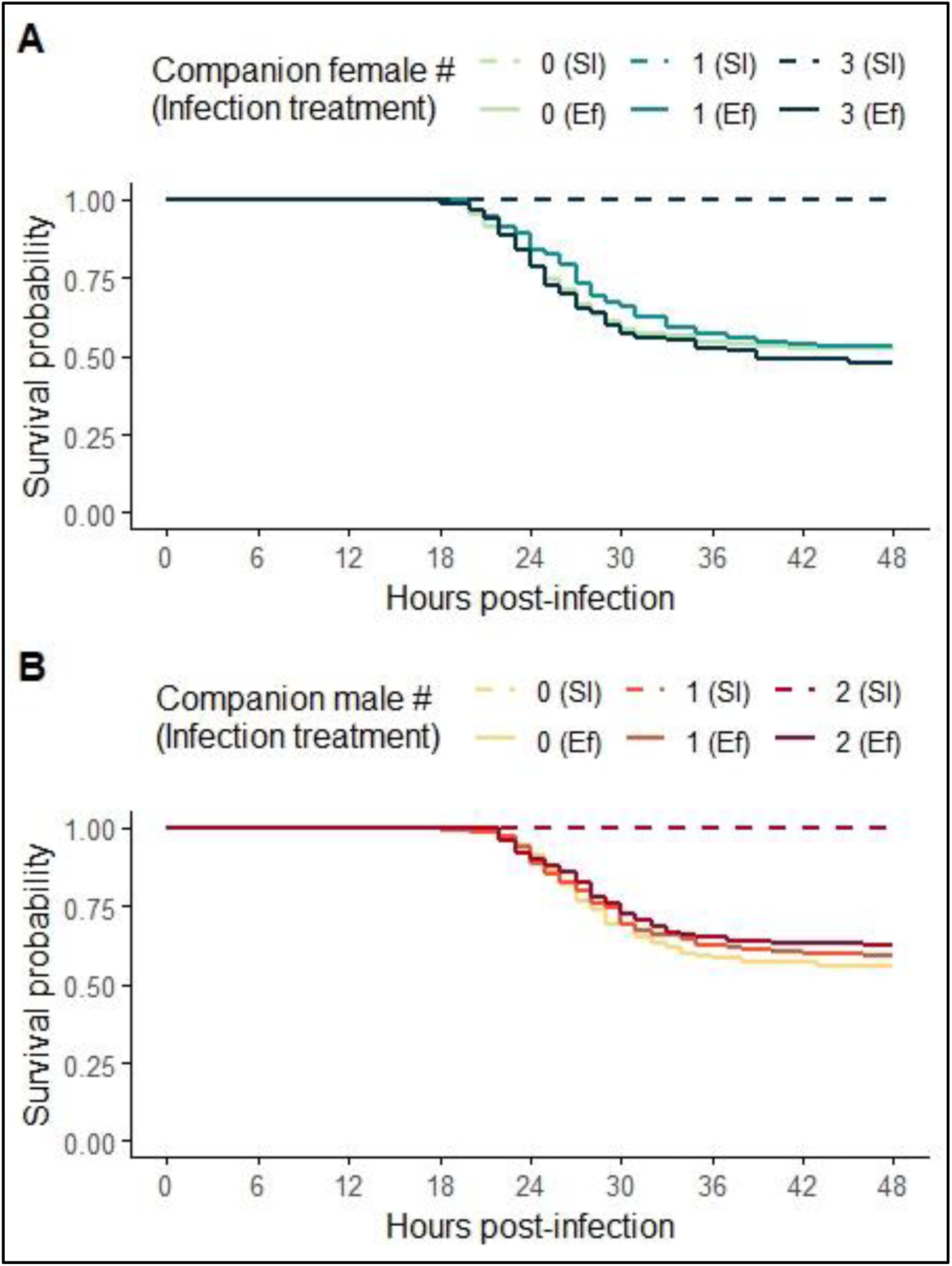
Effect of socio-sexual environment on the survival of *Drosophila melanogaster* females following infection with *Enterococcus faecalis* (Ef) or sham-infection (SI). **(a)** Survival when housed with a varying number of female companions. **(b)** Survival when housed with a varying number of male companions.

This was done to account for the fact that females differ in terms of how long they survive in the infected vs. the sham-infected treatments, and within the infected treatment, and therefore, there is a possibility that a female might produce a greater number of progeny by simply living longer than another female that perished early (Basu et al., 2024a). This *normalized female fecundity was analyzed* using type III Analysis of Variance, with ‘infection treatment’, ‘companion count’, and their interaction as fixed factors and ‘replicate’ as a random factor.

## 3. Results

### 3.1. Experiment 1: Effect of varying numbers of female companions on post-infection survival and fecundity of focal females

Focal females were subjected to infection with *Enterococcus faecalis* or sham infections, and thereafter housed either in isolation, with 1 unmated female, or with 3 unmated females. Post-infection survival and fecundity (number of progeny produced) were measured for each female for 48 hours following infection.

Post-infection survival of females was not affected by the number of female companions (**figure 1(a)**, **table 1(a)**). *Normalised* fecundity (number of progeny divided by female lifespan in hours) was affected by infection treatment, with infected females producing fewer progeny compared to sham-infected females, and by the number of female companions; the interaction between infection treatment and female companion number was not significant (**figure 2(a)**, **table 2(a)**). Housing focal females with 3 companion females decreased fecundity compared to when females were housed alone (**table S1(a)**). There was no difference between the fecundity of focal females housed individually and those housed with 1 companion female, or between the fecundity of focal females housed with 1 companion female or those with 3 companion females (**table S1(a)**).

**Figure 2.**
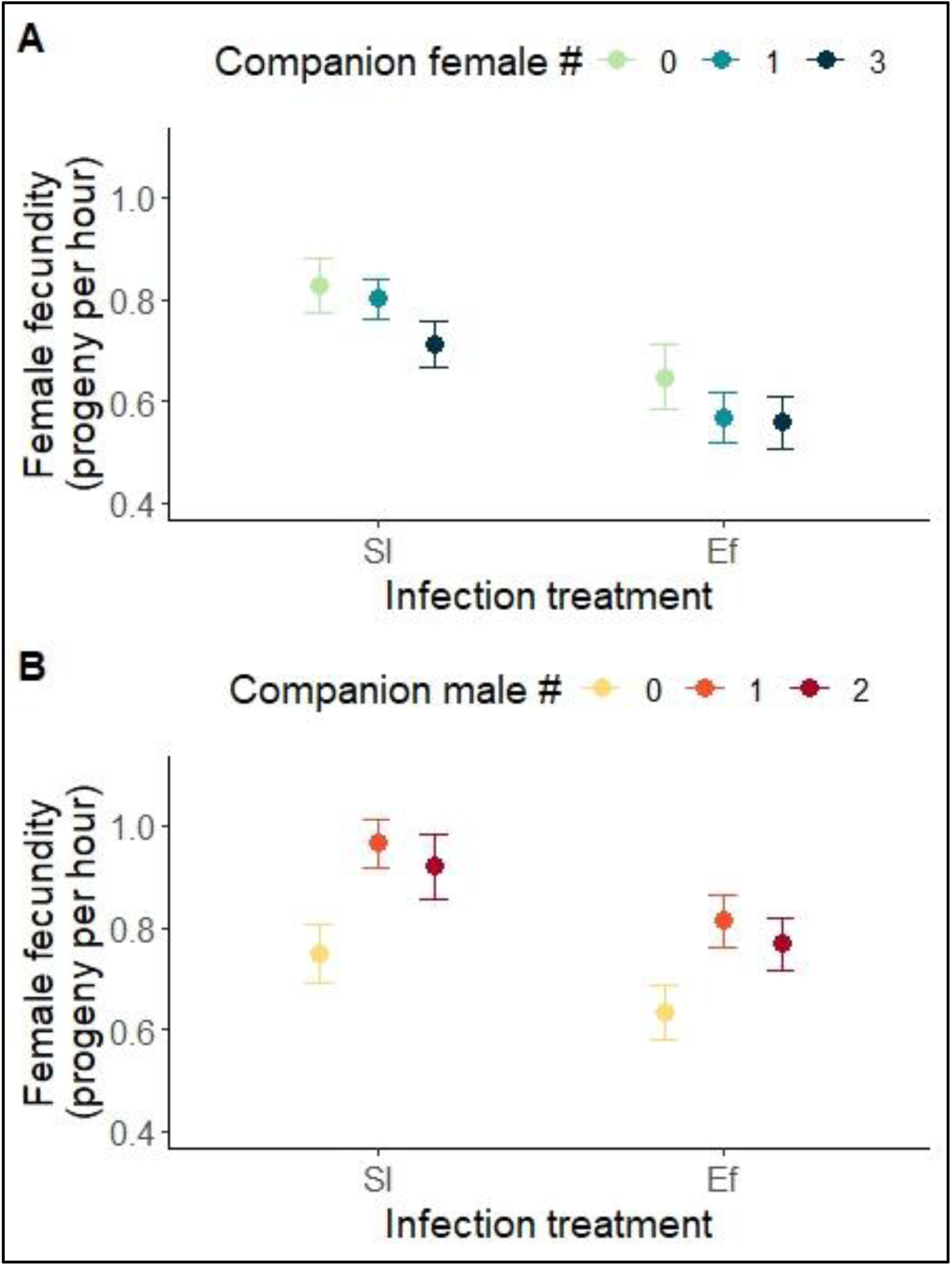
Effect of social and sexual environment on the fecundity of *Drosophila melanogaster* females following infection with *Enterococcus faecalis* (Ef) or sham-infection (SI). **(a)** Fecundity when housed with a varying number of female companions. **(b)** Fecundity when housed with a varying number of male companions.

**Table 1.** Effect of the number of female or male companions on post-infection survival of focal females. Hazard ratios calculated using the Cox proportional-hazards method.

| Treatment | Hazard ratio | Lower CI (95%) | Upper CI (95%) | z value | p value |
| --- | --- | --- | --- | --- | --- |
| **(a) Focal females housed with a different number of companion females** |  |  |  |  |  |
| Isolated focal female (reference treatment) | 1 | na | na | na | na |
| One companion female | 0.907 | 0.671 | 1.225 | -0.64 | 0.52 |
| Three companion females | 1.089 | 0.811 | 1.456 | 0.56 | 0.58 |
| **(b) Focal females housed with a different number of companion males** |  |  |  |  |  |
| Isolated focal female (reference treatment) | 1 | na | na | na | na |
| One companion male | 0.901 | 0.656 | 1.239 | -0.64 | 0.52 |
| Two companion males | 0.830 | 0.600 | 1.149 | -1.12 | 0.26 |

**Table 2.**
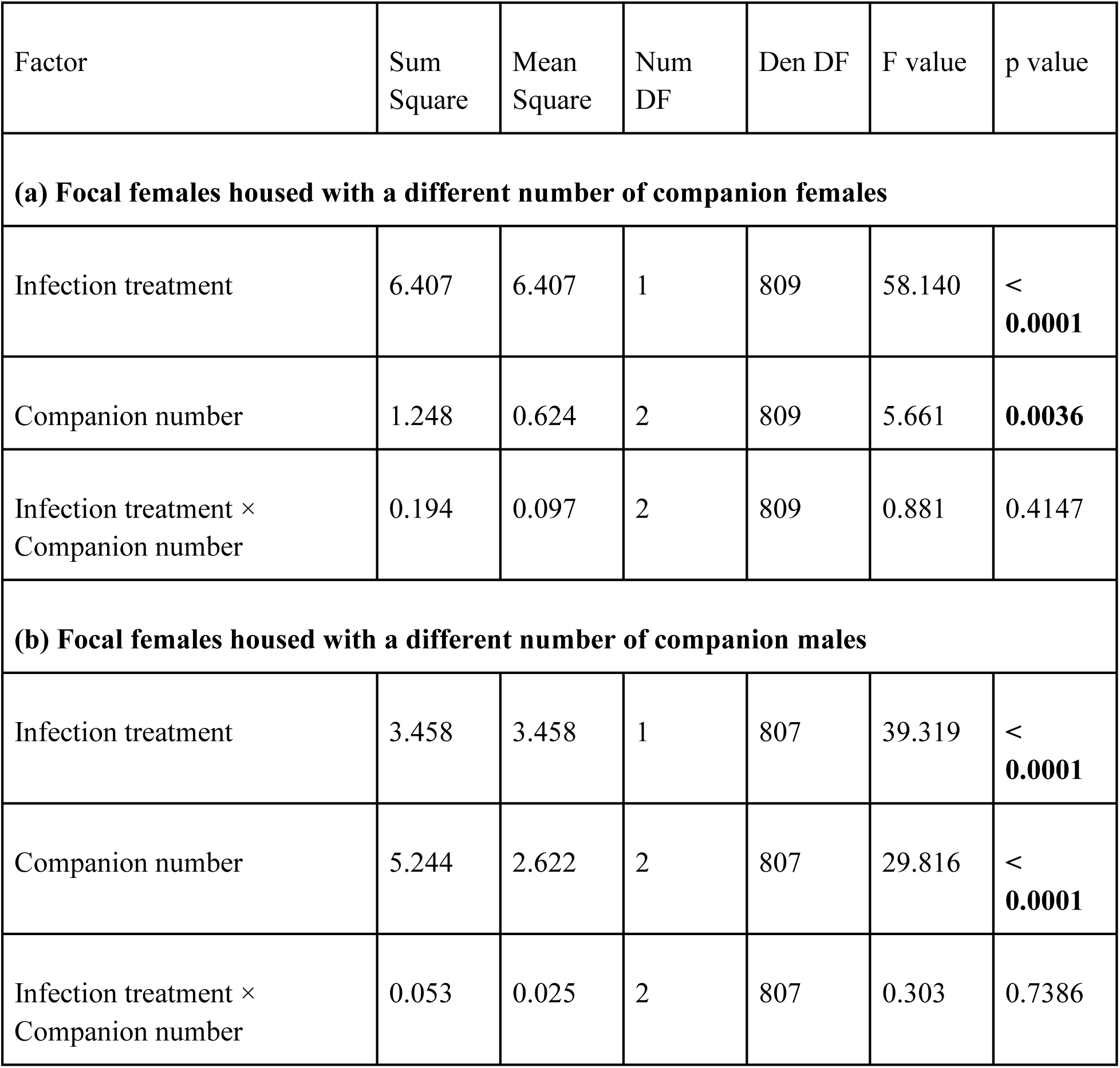
Effect of infection treatment and the number of female or male companions on fecundity (progeny produced per hour) of focal females.

### 3.2. Experiment 2: Effect of varying numbers of male companions on post-infection survival and fecundity of focal females

Focal females were subjected to infection with *E. faecalis* or sham infections, and thereafter housed either in isolation, with 1 male, or with 2 males. Post-infection survival and fecundity (number of progeny produced) were recorded for each female for 48 hours following infection.

Post-infection survival of females was not affected by the number of male companions (**figure 1(b)**, **table 1(b)**). *Normalised* fecundity (number of progeny divided by female lifespan in hours) was affected by infection treatment, with infected females producing fewer progeny compared to sham-infected females, and by the number of male companions; the interaction between infection treatment and male companion number was not significant (**figure 2(b)**, **table 2(b)**). Housing females with 1 male or 2 males increased fecundity compared to when females were housed alone (**table S1(b)**); there was no difference between the fecundity of females housed with 1 or 2 males (**table S1(b)**).

## 4. Discussion

In this study, we investigated whether the socio-sexual environment (SSE) of the host affects the impact of pathogenic bacterial infection on female fecundity in *Drosophila melanogaster*. The SSE of female flies was manipulated by varying the sex and the number of companion flies they were housed with. We housed focal females – either sham-infected (controls), or infected with *Enterococcus faecalis*, a pathogenic bacterium – either in isolation or accompanied by varying numbers of females or males, and measured their post-infection survival and fecundity. Below, we discuss the key results from our experiments and their implications.

### 4.1. Altered SSE alters the fecundity of focal females

Varying the number of companions – females or males – that the focal females were housed with had a significant effect on the fecundity of these females. However, the direction of this effect differed based on the sex of the companion flies. Varying the number of companion females (from 0 to 3) reduced the fecundity of the focal females (**figure 2(a)**), while varying the number of companion males (from 0 to 1 or 2) increased fecundity (**figure 2(b)**).

A reduction in female fecundity with an increasing number of companion females has been previously demonstrated in *D. melanogaster* (Pearl and Parker 1922, Pearl et al., 1926, Pearl 1932, Robertson and Sang 1944a, Churchill et al., 2021, Fowler et al., 2022); however, a few notable nuances are observed in our findings. One, in previous studies, while altering housing densities, authors commonly increased the number of male and female companions simultaneously, and therefore, the effects of female and male companions could not be teased apart. We have demonstrated that female and male companions have distinct effects on female fecundity, with these effects occurring in opposite directions.

Two, in previous studies, the female companions were mated females and therefore competed with one another for oviposition space, in addition to engaging in social interactions (Pearl 1932). Thus, the reduction in female fecundity in those studies was probably driven by the limitation of oviposition space. In our experiments, the companion females were held as virgins, ensuring that they do not compete with focal females over oviposition space. Thus, the reduction in female fecundity we observed is solely driven by social interactions. These differences may also explain why the decrease in fecundity in our experiments is less steep compared to what is reported in previous studies (e.g., Pearl 1932, Robertson and Sang 1944a).

The observed increase in fecundity when the number of companion males was varied was likely due to increased copulation frequency of the focal female in the presence of males (Robertson and Sang 1944b). What is noteworthy here is that although the presence of males in the vial increases female fecundity, increasing the number of companion males from one to two does not lead to a further increase in fecundity.

Overall, these results indicate that the SSE alterations implemented in our experiments were sufficient to induce changes in the reproductive behaviour and physiology of the focal females.

### 4.2. Infection with *Enterococcus faecalis* compromises female fecundity

The focal females infected with *E. faecalis* produced a significantly reduced number of progeny compared to sham-infected females in both of our experiments (**figure 2**). This observation aligns with previous knowledge about this host population and pathogen (Basu et al., 2024a). Reduced fecundity following a pathogenic infection challenge can be interpreted in light of the resource (re)allocation trade-off: increased investment towards anti-bacterial defences can deplete the pool of shared resources, thus reducing fecundity (Sheldon and Verhulst 1996, Lochmiller and Deerenberg 2000, Sadd and Schmid-Hempel 2009). Alternatively, reduced post-infection fecundity may be caused by infection-induced somatic damage, particularly damage to reproductive tissues, resulting from either the pathogen or the host immune response (Brandt and Schneider 2007, Sadd and Siva-Jothy 2006). Our results suggest that the latter scenario is more plausible, since we observe that altering host reproductive effort via altered SSE does not lead to a concomitant change in post-infection survival (also **see sections 4.4 and 4.5**), which would be expected if a resource-allocation-based trade-off were involved.

### 4.3. The effect of infection on fecundity is not contingent on SSE

The primary aim of our experiment was to test whether SSE influences the effect of pathogenic bacterial infection on female fecundity. Contrary to our expectations, SSE did not influence the effect of *E. faecalis* infection on female fecundity (**figure 2**): females, upon being infected, exhibit a similar reduction in progeny production irrespective of whether they are housed individually or in the company of other females or males.

The effect of a pathogenic infection challenge on host fecundity can be contingent on various factors (Duffield et al., 2017). Previous research has demonstrated that the effect of bacterial infections on *D. melanogaster* female fecundity can be pathogen-specific and depends on host genetics, diet, and the phase of infection (Brandt and Schneider 2007, McKean et al., 2008, Linder and Promislow 2009, Howick and Lazzaro 2014, Kutzer and Armitage 2016, Kutzer et al., 2018, Kutzer et al., 2019, Hudson et al., 2020, Basu et al., 2024a). Our results demonstrate that SSE (and the accompanying changes in housing density) does not influence how bacterial infection affects female fecundity in *D. melanogaster*, at least within the parameter space we have explored. Our experimental results may be pathogen-specific, and they do not rule out the possibility that larger manipulations of SSE (and housing density) might lead to different experimental outcomes.

It should be noted that the focal females in our experiments experienced a common housing environment from their eclosion as adults till they were distributed to infection treatments. Any manipulation of the SSE was implemented at the point of infection: that is, the focal females experienced altered SSE from immediately after being infected. Therefore, whatever influence the alteration of SSE had on fly physiology and behaviour was rapid-onset in nature, taking effect in a matter of hours. Hence, it is plausible that long-term manipulation of SSE might have an effect on fly response to infection (in terms of survival and fecundity), even though short-term manipulation, like in our experiments, did not have an effect (e.g., Leech et al., 2019).

### 4.4. Altered SSE does not affect post-infection survival

Post-infection survival of the focal females was not affected by their SSE in our experiments. Housing with or varying the number of either female (**figure 1(a)**) or male (**figure 1(b)**) companions did not make the focal females any more or less susceptible to *E. faecalis* infection than females housed individually. The lack of effect of SSE on post-infection survival is surprising, as altered SSE significantly alters reproductive effort within our experimental parameters; however, it is not entirely unexpected.

Like any study that manipulates the SSE of an organism, our experimental treatments with altered SSE also involved a concomitant change in housing density, regardless of how density is defined (total number of flies around the focal individual vs. the number of flies per unit space or food surface). However, the range of density covered in our experiments was quite small: from females housed individually to females housed in same-sex groups of 4 individuals, housed in the same space with the same access to food. Such small changes in housing density have previously been shown to be inconsequential in terms of post-infection female survival, especially for young females like those used in our study, when infected with multiple different pathogens (Leech et al., 2019). Previous work also suggests that, in the particular host population we used, large changes in housing densities are necessary for survival post-infection with *E. faecalis* to be affected (Das et al., 2022).

In the experiment with varying numbers of male companions, increased immediate reproductive effort, resulting from increased mating frequency in the presence of males, should ideally be interpreted in the context of mate harm (Kuijper et al., 2006), which often manifests as reduced survival (Fowler and Partridge 1989, Chapman and Partridge 1996) but increased short-term fecundity (Hollis et al., 2019). Despite this precedent, we did not observe an effect of male presence on post-infection survival in our second experiment, although a significant increase in fecundity was observed. Our observations align with the existing literature, which suggests that excess mating does not compromise female immune function beyond the effects of a single bout of mating (McKean and Nunney 2005, Gordon et al., 2022). Additionally, this also suggests that mate harm does not necessarily manifest as compromised immune function or increased susceptibility to infection in *D. melanogaster*, at least in the short term, although such an effect has been previously hypothesised (Fedorka and Zuk 2005, Imroze and Prasad 2011).

Additionally, it is noteworthy that although mating is generally known to increase susceptibility of *D. melanogaster* females to bacterial infections, the results are inconsistent between studies for the pathogen we used. One previous study has reported that mated females are more susceptible to *E. faecalis* infections compared to unmated females (Basu et al., 2024c). In contrast, two other independent studies have reported that mating does not affect susceptibility to infections with this pathogen (Short and Lazzaro 2010, Basu et al., 2024b).

### 4.5. Reproduction-immunity trade-off is complex and nonreciprocal

Reproduction–immune function trade-offs are hypothesised to be reciprocal, resource allocation-based trade-offs (Lawniczak et al., 2007, Schwenke et al., 2016). Increased reproductive effort is expected to compromise immune function, thereby reducing post-infection survival, while increased investment in immune defences (brought about by an infection challenge) is expected to adversely affect host fecundity. The reciprocal nature of these trade-offs is rarely tested in a single study (Adhikari and Lazzaro 2025).

In our experiments, infection with *E. faecalis* resulted in a decrease in progeny production, whereas increased progeny production induced by altered SSE did not result in a concomitant change in post-infection survival. Our results thus suggest a lack of reciprocity in reproduction–immune function tradeoffs, and further strengthen our conclusion that the infection-induced fecundity reduction observed in our experiments is due to somatic damage, rather than the reallocation of limiting resources (**see section 4.2**). It is reasonably possible that our results are unique to the host population or the pathogen strain used; however, similar results have been previously reported in *D. melanogaster*, where a change in fecundity due to manipulation of larval diet did not alter susceptibility to *Pseudomonas entomophila* infection (Savola et al., 2022). Similar complex and nonreciprocal manifestation of reproduction–immune function trade-offs are also known in *Tenebrio molitor* (Jehan et al., 2021, Jehan et al., 2022).

### 4.6. Conclusion

We have demonstrated in these experiments that changes in the socio-sexual environment (and accompanying changes in housing density) alter the reproductive effort (measured as the number of progeny produced) in *Drosophila melanogaster* females. Despite this, changes in the socio-sexual environment did not affect post-infection survival or fecundity when females were infected with the bacterial pathogen *Enterococcus faecalis*. Our results thus suggest that the socio-sexual environment does not affect the reproduction-immunity trade-off in *D. melanogaster* females, in that the adverse effect of pathogenic infection challenge on female fecundity was independent of its socio-sexual environment.

---

**Supplementary table S1.** Post-hoc pair-wise comparisons using Tukey’s HSD for the effect of companion numbers on fecundity of focal females.

| Comparison<br>(number of<br>companions) | Estimate | SE | DF | t ratio | p value |
| --- | --- | --- | --- | --- | --- |
| **(a) Focal females housed with a different number of companion females** |  |  |  |  |  |
| Zero vs. One | 0.0534 | 0.0304 | 813 | 1.755 | 0.1857 |
| Zero vs. Three | 0.1020 | 0.0304 | 813 | 3.351 | **0.0024** |
| One vs. Three | 0.0486 | 0.0304 | 813 | 1.597 | 0.2475 |
| **(b) Focal females housed with a different number of companion males** |  |  |  |  |  |
| Zero vs. One | -0.1993 | 0.0272 | 812 | -7.341 | **< 0.0001** |
| Zero vs. Two | -0.1542 | 0.0272 | 812 | -5.678 | **< 0.0001** |
| One vs. Two | 0.0452 | 0.0272 | 812 | 1.663 | 0.2202 |

## Author contributions (CRediT statement)

**Suhaas Sehgal** (Investigation, Data curation, Validation, Writing – review & editing), **Aabeer Kumar Basu** (Conceptualisation, Methodology, Investigation, Data curation, Validation, Formal analysis, Visualisation, Writing – original draft, Writing – review & editing) and **Nagaraj Guru Prasad** (Funding acquisition, Supervision, Writing – review &editing)

## Declaration of generative AI tools

No generative AI tools were utilised during the writing of this manuscript or during any other process concerning the experiments reported in this manuscript.

## Acknowledgements

The authors thank Prof. Brian Lazzaro (Cornell University, USA) for providing the *Enterococcus faecalis* isolate used in the experiments.

